# Genome sequence of *Ophryocystis elektroscirrha*, an apicomplexan parasite of monarch butterflies: cryptic diversity and response to host-sequestered plant chemicals

**DOI:** 10.1101/2023.02.01.526615

**Authors:** Andrew J. Mongue, Simon H. Martin, Rachel E. V. Manweiler, Helena Scullion, Jordyn L. Koehn, Jacobus C. de Roode, James R. Walters

## Abstract

Apicomplexa are ancient and diverse organisms which have been poorly characterized by modern genomics. To better understand the evolution and diversity of these single-celled eukaryotes, we sequenced the genome of *Ophryocystis elektroscirrha*, a parasite of monarch butterflies, *Danaus plexippus*. The genome is miniscule, totaling only 9 million bases and containing fewer than 3,000 genes. We then compared this new sequence to the two other sequenced invertebrate-infecting apicomplexans, *Porospora gigantea* and *Gregarina niphandrodes*, which have nearly twice the gene content and found that *O. elektroscirrha* shares different orthologs with each sequenced relative, suggesting the true set of universally conserved apicomplexan genes is very small indeed. We investigated sequenced reads from other potential hosts to explore the viability of *in silico* infection screening. We recovered a similarly sized parasite genome from another butterfly, *Danaus chrysippus*, that was highly diverged from the *O. elektroscirrha* reference, possibly representing a distinct species. Using these two new genomes, we investigated potential evolutionary response by parasites to toxic phytochemicals their hosts ingest and sequester. Monarch butterflies are well-known to tolerate toxic cardenolides thanks to changes in the sequence of their Type II ATPase sodium pumps. We show that *Ophryocystis* completely lacks Type II or Type 4 sodium pumps, and related proteins PMCA calcium pumps show extreme sequence divergence compared to other Apicomplexa, demonstrating new avenues of research opened by genome sequencing of non-model Apicomplexa.

**Author Summary:** There are many relatives of *Plasmodium*, the single-celled parasites responsible for malaria, and they infect a wide range of animals, including insects. These parasites have received less attention however, leaving much unknown about them. We sequenced the genome of one such parasite, *Ophryocystis elektroscirrha* (OE), to compare it to better-studied parasites and shed light on OE’s interaction with its host, the monarch butterfly. We found that OE has a tiny genome with the fewest genes of any sequenced parasite in this group, lacking many genes found in its relatives. Using our new data, we also discovered evidence that infections of other butterfly species that have been attributed to OE may be caused by a previously undiscovered distinct parasite species. And finally, we explored the evolution of a family of genes that may be targeted by medicinal plant compounds in the monarch butterfly’s diet; OE has lost one of these genes and radically changed the sequence of another, setting a direction for future research.

## Introduction

Eukaryotic genomics has overwhelmingly focused on multicellular organisms (1), in spite of the staggering diversity of unicellular eukaryotes. Furthermore, although animal genome sequencing efforts are increasingly aiming to systematically characterize extant species’ genomes (2–4), when unicellular eukaryotes are studied, it is still mostly in connection to human health or economic interests; in other words, by putting most of the effort into studying pathogens of human-importance, the scientific community has left large gaps in our knowledge of these ubiquitous organisms.

One of the most prominent examples of this deficiency is the Apicomplexa (5). This phylum in the Kingdom Protista consists entirely of single-celled parasites and includes the organisms responsible for human diseases such as malaria, caused by *Plasmodium spp*., and toxoplasmosis, from infections with *Toxoplasma gondii* (6). Much of the genetic study of Apicomplexa has been directed at these and similar taxa. However, apicomplexans parasitize other vertebrates (see Levine 1986 for numerous examples) as well as a wide range of invertebrates (8,9). There have been far fewer studies of the latter group of parasites and their diversity largely remains unexplored with morphological, let alone genomic methods (10).

With the rise of genome sequencing, many such apicomplexans are identified incidentally in sequencing data generated from infected hosts (11), though there are problems with this passive sampling approach. Oftentimes, these contaminants are not apparent in the assembly process owing to both the poor taxonomic sampling of Apicomplexa and the miniscule size of their genomes. Limited taxonomic representation in available databases means that contamination may not be detected via sequence similarity searches and any outlier sequences, e.g. differing in from the host in GC content, may not represent a large enough proportion of an assembly to merit further investigation. The largest genomes of sequenced species, *Toxoplasma gondii* and *Neospora canum*, are only 70 megabases in length (12,13), and many species have genomes totaling less than 10 megabases (14,15). With metazoan genomes typically ranging from hundreds of megabases to several gigabases, these Apicomplexan genomes are all < 10% but typically closer to 1% the size of their hosts’ genomes. Note, however, that these genome size metrics come from vertebrate-infecting Apicomplexa, limiting the generalizability of these patterns. Here we sequence and annotate the genome of an insect pathogenic apicomplexan with a wealth of ecological data but a paucity of molecular resources: *Ophryocystis elektroscirrha*.

### Ophryocystis elektroscirrha *as a model non-model Apicomplexan parasite*

Among the invertebrate-infecting Apicomplexa, one of the best-studied is the neogregarine *Ophryocystis elektroscirrha* (Neogregarinorida: Ophryocystidae, McLaughlin and Myers 1970). It parasitizes monarch butterflies, *Danaus plexippus* (Lepidoptera: Nymphalidae), beginning when caterpillars ingest oocysts shed onto eggs and milkweed plant matter by infected adult butterflies. Within the host’s gut, the oocyst opens, releasing eight parasite cells that invade the caterpillar’s tissues and then asexually reproduce, creating clones of the initial infectious genotypes. During the host’s pupal stage, the parasites sexually reproduce to form new oocysts with two copies of four meiotic products, for a total of eight cells per oocyst. These oocysts form on the cuticle of the developing butterfly, ultimately residing on scales which easily fall onto eggs and plants as the adult butterfly feeds and lays eggs (8). Infected monarchs have shortened lifespans and decreased flight performance; those with the heaviest infections often fail to emerge from the pupa and quickly die (16,17). These costs interact with the unique behavioral ecology of monarchs to create a complex but well-characterized host-parasite system.

Most North American monarchs belong to one panmictic population that annually migrates dozens to thousands of kilometers to overwinter in warmer climates (18). Some monarch populations, however, for example in southern Florida, experience stable climate and milkweed availability year-round and do not migrate; these non-migratory populations show much higher prevalence for *O. elektroscirrha* infection than their migratory counterparts (19,20). These observations suggest that migratory behavior limits parasite prevalence, because infected individuals are less likely to complete the journey than their uninfected counterparts (21,22). In addition, climate conditions that induce migration in the butterflies also cause senescence of native milkweed hostplants. Because caterpillars become infected by consuming oocysts shed onto eggs and these plants by adults, host plants can become reservoirs for disease as the season progresses. Indeed, monarch butterflies that associate with persistent perennial milkweed plants are more likely to be infected than their migratory counterparts (23).

The milkweed plants introduce another layer of interaction in the host-parasite system. Monarchs feed on several species of milkweed in the genus *Asclepias* which vary in the levels of phytochemicals they produce. Milkweeds produce cardiac glycosides called cardenolides, which are toxic to vertebrate predators (24) but sequestered by monarch caterpillars as a chemical defense (25). Monarchs themselves remain largely unaffected thanks to a small set of mutations in their Na^+^ /K^+^ – ATPase (sodium potassium pump) enzymes (26). Intriguingly, parasite infection and virulence is strongly affected by plant chemistry; more toxic milkweeds, with more cardenolides, more strongly inhibit parasite growth (27). Even indirect effects, such as the presence of secondary milkweed herbivores, have demonstrated effects on *O. elektroscirrha* infection dynamics in relation to plant chemistry (28). In summary, the dynamics of this butterfly-parasite relationship have been well studied from the local to continental scale. However, in both the case of migration and response to milkweed chemistry, a fundamental research imbalance is evident: the genetics of the monarch butterfly are well characterized, but those of *O. elektroscirrha* remain unexplored, leaving basic questions unanswered. How does parasite population structure compare to that of the host? And is there evolution of *O. elektroscirrha* in response to milkweed chemistry? To begin to answer those questions, we present the genome sequence of *Ophryocystis elektroscirrha*.

First, we apply this new resource to the question of apicomplexan diversity and host specificity. The initial description of this parasite described both *Danaus plexippus* and *Danaus gilippus* as hosts (8). Since then, similar apicomplexan infections have been reported in other members of the *Danaus* genus, including *D. eresimus, D. petilia* (29), and *D. chrysippus* (30). Even apicomplexan infections of the more distantly related butterfly *Parthenos sylvia* and of moths in the genus *Helicoverpa* have been attributed to *O. elektroscirrha*-like parasites(31). These diagnoses are based mainly on the gross morphological similarity of oocysts and may be limited by a paucity of informative morphological characters. As such, it is presently unclear whether *O. elektroscirrha* is a generalist lepidopteran parasite or if there is unrecognized diversity between host species. In addition to characterization of the genome, we demonstrate its use in identifying a likely cryptic species found in sequences from a related butterfly, *Danaus chrysippus*.

Second, we analyze the newly generated sequences to examine the potential for compensatory evolution to resist milkweed cardiac glycosides. It has already been demonstrated that other milkweed-feeding insects evolved parallel amino acid changes in Na^+^ /K^+^ ATPases to tolerate these chemicals (32,33). Moreover, these insects exist within ecological communities of predators and parasites that feed upon them, and a number of birds and parasitoid insects likewise have converged on similar resistant genotypes (34). ^+^

Apicomplexan biochemistry is generally less well understood than that of Metazoa. Indeed, Apicomplexa were long thought to lack Na^+^ ATPases, instead relying solely on Ca^2+^ ATPases, also known as calcium pumps (35). However, it has recently been shown that this initial functional annotation (and likely comparative annotations relying on it) were incorrect, and many Apicomplexa do in fact possess distinct Na ATPases (36,37). We use the new *Ophryocystis* genomes to explore ATPase gene family evolution in Apicomplexa and their potential relationship to cardiac glycosides.

## Methods

### Parasite growth and propagation

We initially collected *O. elektroscirrha* from wild-caught eastern migratory monarch butterflies (strain E41-1a) and propagated them in a laboratory setting; we fed second instar caterpillars a leaf disk containing a single oocyst to establish infections (following previously designed infection methods: 27,38). When the infected adult butterflies eclosed from their pupae, we froze them before collection of oocysts from the outsides of their bodies. Because oocysts are the result of meiotic cell division, this method results in a mix of related parasite genotypes rather than strictly identical clones but is ultimately the only way to generate enough parasite cells for DNA extraction and sequencing.

### Concentration and purification of oocysts

We removed wings from the infected butterfly bodies and vortexed the bodies for 5 minutes in 100% ethanol in glass scintillation vials, a modification of de Roode et al. (38). Oocysts, like the scales with which they associate, entered solution better in ethanol than water, generating a mix of both parasite oocysts and host scales. To separate scales from oocysts, we passed the solution through a 30μm cell straining filter (Miltenyi Biotec, Bergisch Gladbach, Germany), which captured the scales (>100 μm length) while allowing the oocysts to pass through. We centrifuged the flow-through at 14,000 x g for 2 minutes to pellet the parasite oocysts and combined pellets across butterfly hosts to increase yield. Ultimately, we used oocysts from eight butterflies infected with the same initial parasite isolate (i.e. different oocysts from the same initial infected host) for DNA extraction.

### Extraction and sequencing

The thick protein shell of the oocysts strongly inhibited lysing of parasite cells to access DNA. Thus, we needed to physically disrupt the oocysts prior to extraction. We ground the pellet with a Dounce homogenizer for 1.5 hours in lysis buffer on ice, examining an aliquot under a light microscope every 15 minutes and stopped after nearly all of oocysts were visibly broken. Note that oocyst disruption began long before the 1-hour mark, so molecular protocols with lower input DNA requirements (e.g. PCR-based assays) would likely find success even with a greatly reduced disruption phase of this protocol. After this lengthy homogenization step, we followed the kit standard protocol for Omniprep extraction of genomic DNA (G-Biosciences, St. Louis, MO) and sequenced 250 basepair paired-end reads on an Illumina MiSeq.

To aid in assembly and annotation, we also generated RNA sequencing for *O. elektroscirrha* by extracting RNA from a heavily infected monarch pupa. We chose this host-stage for three reasons. First, *O. elektroscirrha* migrates to the cuticle and undergoes oocyst formation at this stage, guaranteeing that the parasite is transcriptionally active. Second, as this is the final active stage before going dormant in oocysts, it represents the peak number of parasite cells in the host. Finally, the level of infection is easily visible as dark spots on the green butterfly pupa (17), allowing us to manually select the most infected individual for extraction. Although contamination with host tissue and transcripts is practically unavoidable at this stage, we aimed to minimize contaminants by using dissecting scissors to target the dark aggregations of *O. elektroscirrha* while avoiding less infected host tissue. We extracted RNA using a Qiagen RNAeasy extraction kit (Hilden, Germany) and carried out Illumina 100bp paired-end sequencing.

### Read processing and assembly

We were concerned that the tight association between host and parasite could still lead to accidental sequencing of host DNA in spite of our upstream attempts to separate the two tissue sources. As a final precaution against contamination, we first aligned raw sequenced reads to the monarch reference genome (39) with bowtie2 using the very-sensitive-local alignment algorithm (40) and only kept unmapped reads for assembly. This methodology may weaken the power to detect horizontally transferred genes with conserved sequence but minimizes the chances of erroneously incorporating host sequence into the parasite assembly. With these high-confidence parasite reads, we sought first to set expectations prior to assembly using k-mer based methods. We counted k-mers using Jellyfish v2.2.6 (41) and then plotted the resultant k-mer frequency histogram and used this distribution to estimate genome size using custom scripts in R v3.3.3 (42). Finally, we assembled these filtered reads with SPAdes v3.13 (43), using a k-mer coverage cutoff of 100 based on the characterization from the previous step. All other parameters were default settings for the tool.

### Scaffolding

We aligned the RNAseq dataset to the newly generated assembly using TopHat v2.1.1 (44) then used Rascaf v1.0.2 (45) to scaffold the assembly based on RNA alignment. We assessed improvement to assembly summary statistics (N50 and contiguity) with QUAST v4.6.3 (46). To assess the improvement in assembly of coding regions, we took both the original and scaffolded assembly through the annotation process described below, ending in evaluation with BUSCO v5.0 (47).

### Annotation

We annotated the initial and RNA-scaffolded assemblies using the GenSAS web-based pipeline (48). Prior to uploading, we trimmed the assemblies to remove contigs totaling fewer than 2,500 bases in length, both to meet requirements of the pipeline and because these contigs were unlikely to contain complete gene sequences. This approach resembles a recently successful effort to characterize the apicomplexan *Porospora gigantea* (49). We then soft-masked the trimmed assembly after two independent rounds of repeat annotation, first with RepeatMasker v4.1.1 and then with RepeatModeler v2.0.1 before combining the two to mask bases for downstream analysis.

We employed two approaches to gene modeling, namely, BRAKER v2.1.1 (50) using alignment of RNA sequencing from the infected host pupa to the draft parasite genome, and GeneMark-ES v4.48 *ab initio* (51). We evaluated both of these predicted gene sets using BUSCO v5.0 to search against the apicomplexa_odb10 database (47) and selected the gene set that maximized the number of identified single copy orthologs to use as the official gene set. Finally, to contextualize this new annotation with existing genome assemblies, we compared conserved orthologs against other gregarine Apicomplexa, namely the annotated protein sequences of *Gregarina niphandrodes* (unpublished, but available via the bioproject: PRJNA259233) and the two recently released *Porospora* genomes (49). Again we used BUSCO v5.0’s apicomplexa_odb10 database (47).

### Screening for infection in genome resequencing data

With the newly generated genome, we explored the potential to detect infection by *Ophryocystis* in genome resequencing data generated from butterfly tissues. We used published Illumina resequencing datasets (100 or 150 bp paired-end) from seven species (Zhan *et al*. 2014; Martin *et al*. 2020, see supplemental files for sample details and accession numbers). We aligned Illumina reads to the masked *O. elektroscirrha* genome using bwa MEM version 0.7.17 (54) with default parameters. We used SAMtools version 1.9 (55) for conversion of SAM to BAM format, and Picard version 2.21.1 (56) SortSam and MarkDuplicates for sorting and removal of PCR duplicate reads. Given the massive overrepresentation of host butterfly DNA in the data, we took precautions to minimize misalignment of host DNA to the parasite genome. Specifically, only reads with a mapping quality of at least 60 were retained in the SAMtools step, and we subsequently removed all alignments of less than 100bp in length, ignoring indels and soft clipping, using a custom Python script. We then computed mean read depth per scaffold using Mosdepth (57).

### Screening for infection by detection of oocysts

For 18 *D. chrysippus* samples (Figure S1 and supplemental data tables) we had both Illumina resequencing data and preserved butterfly bodies, allowing us to compare the detection of infection using genomic data with the conventional approach of visually screening for oocysts under a microscope. Because the bodies were preserved in ethanol, we used a modified diagnostic procedure: A pipette was pressed against the abdomen and 10 µl of ethanol mixed with scales was pipetted onto a clean microscope slide, which was then viewed under 10x and 40x magnification for detection of oocysts. If too few scales were present on the slide, the procedure was repeated.

### *Identifying* Ophryocystis *scaffolds in a* Danaus chrysippus *assembly*

Given that a *D. chrysippus* pupa used for a previous genome assembly (53) showed evidence for infection by *Ophryocystis* (see Results), we attempted to identify *Ophryocystis* scaffolds in the *D. chrysippus* assembly. We used a version of the assembly prior to removal of small scaffolds to ensure maximal recovery. We first generated a whole genome alignment between the *D. chrysippus* and *O. elektroscirrha* assemblies using minimap2 (58), with the ‘asm20’ parameter present, which is optimized for more dissimilar genomes. We then removed alignments less than 100 bp in length and those with sequence divergence (dv tag output by minimap2) >0.15, based on visual inspection of the divergence distribution. Given the availability of a new highly complete *D. chrysippus* assembly from an uninfected individual (59), we generated a second whole genome alignment between the infected and uninfected *D. chrysippus* assemblies using the same procedure and filters. We reasoned that scaffolds representing *Ophryocystis* in the infected assembly should have strong homology to the *O. elektroscirrha* genome and little or no homology to the uninfected *D. chrysippus* genome. However, given that repetitive sequences such as transposable elements (TEs) may be shared between host and parasite, we do not expect a complete lack of homology between parasite and host scaffolds. Based on visual inspection of the data, scaffolds were defined as confidently representing *Ophryocystis* if (1) alignments to *O. elektroscirrha* comprised at least half the scaffold length (after accounting for overlapping alignments), and (2) alignments to the uninfected *D. chrysippus* comprised less than a third of the scaffold length aligned to *O. elektroscirrha*.

As an additional line of evidence to exclude false positives, we considered read depth of Illumina reads from one infected and one uninfected adult butterfly (detected as described above). We reasoned that scaffolds representing *Ophryocystis* should have non-zero average read depth for reads from the infected adult, but zero read depth for reads from the uninfected adult. Given the low read depths (see Results) we required that scaffolds had a mean depth of at least 0.1 for the infected butterfly. Due to possible mis-mapping and shared repetitive sequences such as TEs, we relaxed the expectation that normalized read depth should be zero in the uninfected butterfly, and instead required that it must be lower than that in the infected butterfly.

Finally, we took these apicomplexan sequences through the same gene annotation pipeline as we used for our de novo *O. elektroscirrha* assembly. Ultimately, we compared the number and identity of BUSCO orthologs identified in each. Note however that this *D. chrysippus*-derived assembly is less contiguous owing to its incidental assembly within the host genome, so we discount cases of apparently missing orthologs as potential false negatives.

### Phylogenetic diversity of Ophryocystis

The ability to extract *Ophryocystis* sequence reads from genomic data of infected butterflies allowed us to investigate *Ophryocystis* diversity using previously published sequence data. We selected the twelve samples with the highest mean read depth for this purpose. Using the same filtered BAM files described above, we called haploid genotypes using bcftools v1.10.2 (55) using the mpileup, call and filter tools to retain only non-indel genotypes with genotype quality (GQ) ≥20. Due to low sequencing depth (<1x for most individuals) we could not reliably filter out potentially repetitive regions based on individual read depth. Instead, we excluded regions where the total read depth across all 12 individuals was > 30x, based on visual inspection of the genome-wide distribution. Our use of haploid genotypes assumes that each butterfly is infected by only a single parasite strain. Violation of this assumption would lead to a phylogenetic analysis in which each tip represents a combination of ancestries, which is in fact true for all genome-scale phylogenetics of recombining species. We therefore chose to use the distance-based neighbor-joining method, which is a genetic clustering algorithm that does not rely on an underlying model of a bifurcating tree. Pairwise genetic distances were computed using the distMat.py script https://github.com/simonhmartin/genomics_general), considering only sites with genotypes for at least 9 of the 12 infected individuals. A neighbor-joining tree was generated using the BIONJ method implemented in splitsTree v4 (60).

### Investigation of parasite ATPase evolution

We searched for signatures of co-evolutionary dynamics in the gene content of *Ophryocystis* in relation to its host biochemical environment. In particular, it has been well documented that *Danaus plexippus’s* cardiac glycoside-rich diet has driven changes in the amino acid sequence of its Na^+^ /K ^+^ – ATPase, making them tolerant of the effects of this toxin (26,32,33). Given that *Ophryocystis* spends part of its life cycle in the host’s gut, surrounded by cardiac glycosides and is negatively impacted by these toxins (27), it is reasonable to assume this interaction exerts selective pressure on ATPases in the parasite as well. Even in the relatively reduced gene sets of Apicomplexa, many species have multiple ATPases which pump different cations and fulfill different cellular roles. To explore ATPase dynamics in these parasites required us to both identify relevant sequences and infer their putative functions. ^+^

To start, we obtained the protein sequence of the *Plasmodium falciparum* sodium potassium ATPase (accession AAF17245). This ATPase, a P-type ATPase 4, is often labeled as a Ca^2+^ ion pump, based on previous mis-annotation, but is likely an Na – ATPase as shown by more recent work in *Toxoplasma* (35–37). We used BLAST+ (blastp -evalue 1e-10; Camacho *et al*. 2009) to extract homologous genes from the amino acid sequences of our newly generated *Ophryocystis* annotations as well those of *Gregarina niphandrodes* and *Porospora gigantea*-A. We then used a phylogenetic approach to place these unannotated gregarine sequences in the context of a recent and more comprehensive dataset of ATPases across Apicomplexa and Metazoa to infer ATPase type and function via sequence similarity (36). We first aligned the gregarine protein sequences using MAFFT (as implemented in the Geneious software (62)), then manually aligned these sequences with the 265 amino acids corresponding to the highly conserved domains among ATPases analyzed by Lehane et al. (36), and finally trimmed the gregarine sequences to contain only those sites corresponding to the conserved 265 amino acids. Using this combined alignment of gregarine and other ATPases, we estimated a maximum likelihood phylogeny using the phangorn package in R, employing the LG+G(4)+I substitution model and 100 bootstrap replicates (63). Subsequently, functional annotations were assigned to the gregarine sequences based on their placements within clades of distinct ATPases, as assessed by Lehane et al. functions assigned Finally we used phylogenetic relationships to infer functional classes of our uncharacterized gregarine ATPases in relation to established ATPases from other species. **Results**

### Raw sequence data composition

Initial alignment showed 28% of total reads aligning to the host genome. We treated the former as butterfly sequences and the remaining 72% of unmapped reads as putative parasite sequences. K-mer analyses with Jellyfish (41) suggested very little heterozygosity and a homozygous read depth of roughly 530x for these unmapped reads. Using this k-mer distribution and custom R scripts, we calculated the expected genome size to be roughly 8.11Mb, which would place *O. elektroscirrha* among the smallest sequenced apicomplexan genomes.

### Assembly summary

An overview of the assembly is available in Table 1 (middle column). Initially, we assembled close to, but more than our k-mer based expectations: 8,864,135 bases. The assembly was contained in 909 contigs, with a minimum size of 128 basepairs and an N50 of 57,260 bases. We also noted that *O. elektroscirrha* is very GC poor (∼27%), a feature characteristic of some other apicomplexan genomes. Before continuing with analyses, we needed to trim the smaller contigs from the assembly to meet criteria for annotation tools, so we set a minimum contig length threshold of 2,500 bases. This trimmed assembly totals 8,629,789 bases in 244 contigs, with an N50 of 58,409 bases. The loss of 234Kb of sequence (roughly 2.6% of the assembly) is a calculated tradeoff to format the genome for further analyses. Moreover, given the extremely high coverage and the method of extracting DNA from oocysts after sexual reproduction, it is likely that many of the very small contigs represent either read errors or regions of high genetic diversity that have enough coverage to not be discarded, but owing to their variation could not be collapsed into single sequences during assembly.

**Table 1.** At-a-glance assembly and annotation statistics for the *Ophryocystis* sequences analyzed here. Raw size denotes the total length of bases *de novo* assembled or extracted from a host assembly in the case of the *O. elektroscirrha*-like assembly from *D. chrysippus*. We discarded sequences fewer than 2,500 bases in length prior to downstream gene annotation, yielding the analyzed size of each genome. We scaffolded the *O. elektroscirrha* assembly with RNA sequencing but did not perform comparable scaffolding for the *O. elektroscirrha*-like assembly due to divergence of sequences and lack of appropriate RNA data. We ran BUSCO on annotations to identify putative universally conserved single copy orthologs across Apicomplexa. Results follow the format: C – complete [S – single-copy, D – duplicated] F – fragmented, M – missing.

|  | *Ophryocystis elektroscirrha* | *O. elektroscirrha*-like |
| --- | --- | --- |
| Raw size | 8,864,135 bp | 8,799,632 bp |
| GC content | 27.18 % | 26.93 % |
| Analyzed size | 8,631,036 bp | 8,058,112 bp |
| Contigs | 909 | 332 |
| Contig N50 | 57,260 bp | 49,155 |
| Scaffolds | 156 | – |
| Scaffold N50 | 89,860 bp | – |
| Repeat content | 11.76% | 11.49 % |
| Annotated genes<br>(protein coding) | 2,633<br>(2,591) | 2,369<br>(2,280) |
| BUSCO apicomplexaodb10<br>(n = 446) | C:80.9% [S:80.5%,D:0.4%],<br>F:1.1%, M:18.0% | C:48.9% [S:48.0%,D:0.9%],<br>F:11.2%, M:39.9% |

Finally, prior to annotation, we scaffolded the trimmed assembly using the RNA-seq data from an infected host pupa. *Ophryocystis elektroscirrha* transcripts were a minority of sequences, as evidenced by an alignment rate of only roughly 10% of sequenced reads. Still, these reads proved sufficient to improve the assembly. After scaffolding with Rascaf, the final assembly consists of 156 scaffolds with an N50 of 89,860 bases. This scaffolding also improved the annotation of genes, as evidenced by an increase in identified BUSCOs reported below.

### Repeat content

As expected of a very small genome, very little of the sequence was repetitive. In all, 1,014,979 bases (11.8% of the trimmed length), were masked. The majority of these, ∼600kb, were unclassified repeats, with the remaining ∼400kb being simple repeats.

### Gene content

We employed two approaches to gene annotation, first using BRAKER to incorporate information from the alignment of RNAseq from an infected host to the *O. elektroscirrha* genome before RNA-scaffolding. With this method, we annotated 2,915 genes (encoding 3,122 proteins). This gene set contained 72.2% of the apicomplexa_odb10 BUSCO genes in a complete state, with a further 6% identifiable as fragmented. We also carried out *ab initio* annotation using GeneMark-ES without evidence beyond the genome sequence itself. We annotated 2,695 genes, 2,632 of which encoded a protein. This gene set contained far more BUSCOs, with 79.1% as complete genes and another 2% fragmented. As this method gave much better BUSCO results than BRAKER, we used it as the standard for subsequent annotations.

First, we reannotated the genome after scaffolding with RNAseq. In this assembly, GeneMark annotated 2,633 genes (2,591 protein coding). Note that although the total number of genes and the number of protein coding genes both decreased, the difference between the two decreased as well, as expected if RNA-scaffolding stitched together previously fragmented genes into complete coding sequences. This gene set contained 80.9% complete and 1.1% fragmented BUSCOs.

To contextualize this new assembly relative to previously studied Apicomplexa, we present summaries of genome size and gene content in both tabular (Table 2) and graphical form (Figure 1). Additionally, we compared conserved gene content between *O. elektroscirrha, P. gigantea*-A, and *G. niphandrodes* (Figure 2). In total, 309 of the 446 (69%) of BUSCO orthologs were conserved in all 3 genomes, with an additional 80 (18%) identifiable in two of three species. A further 32 orthologs (7%) were found only in one species, leaving 25 (6%) absent from all three. Despite *O. elektroscirrha* possessing roughly half the number of genes as the other two gregarines, its gene set is not a simple subset of either. Taken together this suggests substantial lineage-specific gene loss.

**Table 2.** *Ophryocystis elektroscirrha* in the context of other Apicomplexa. Genome size, annotated gene count, and gene density for a selection of published apicomplexan species. *Ophryocystis elektroscirrha* has a smaller genome than most other species and contains the fewest protein-coding genes yet-described for an Apicomplexan. We have excluded the *O. elektroscirrha*-like assembly from this comparison, as our methods of identification and filtering for annotation make it more likely to be an incomplete sequence and annotation.

| Species | Order | Genome size (Mb) | Gene content | Gene density (genes/Mb) | Reference |
| --- | --- | --- | --- | --- | --- |
| *Toxoplasma gondii* | Eucoccidiorida | 61.6 | 8,789 | 142.7 | Yucesan et al., 2021 |
| *Neospora caninum* | Eucoccidiorida | 61.5 | 7,540 | 122.6 | Berná et al., 2021 |
| *Plasmodium falciparum* | Haemospororida | 22.9 | 5,268 | 245.8 | Gardner et al., 2002 |
| *Gregarina niphandrodes* | Eugregarinorida | 14 | 6,375 | 455.4 | Unpublished, bioproject: PRJNA259233 |
| *Cryptosporidium parvum* | Eucoccidiorida | 9.2 | 3,870 | 425.3 | Abrahamsen et al., 2004 |
| *Theileria orientalis* | Piroplasmida | 9.0 | 4,002 | 444.7 | Hayashida et al., 2012 |
| *Ophryocystis elektroscirrha* | Neogregarinorida | 8.8 | 2,633 | 299.2 | This article |
| *Porospora gigantea* | Eugregarinorida | 8.8 | 5,270 | 598.9 | Boisard et al., 2021 |

**Figure 1.**
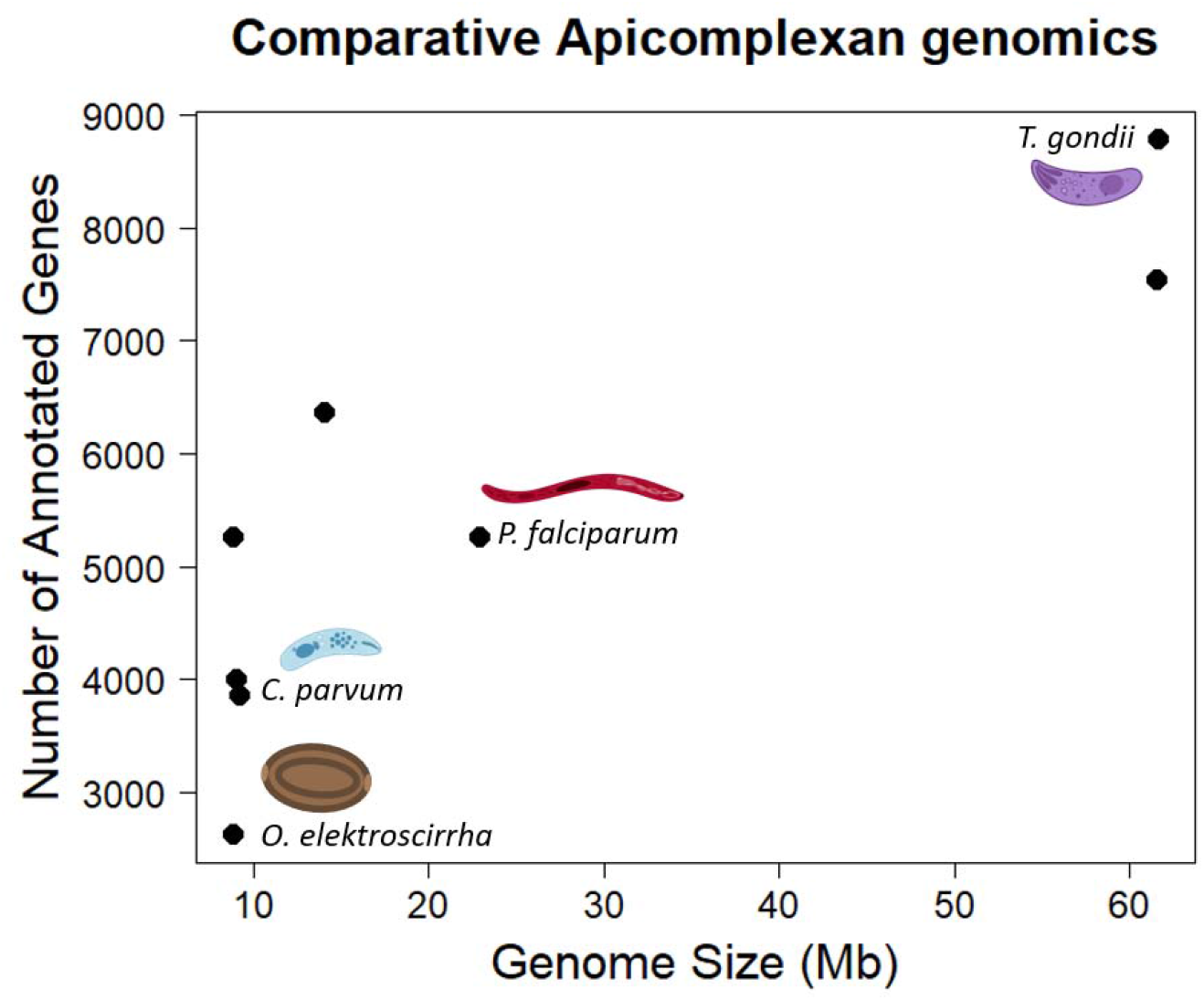
A visual representation of genome size and gene content from Table 2, with notable taxa named and illustrated. Illustrations, except for *O. elektroscirrha* were created with BioRender.com. *Ophryocystis elektroscirrha* has the fewest annotated genes and one of the smallest overall genomes yet sequenced in Apicomplexa.

**Figure 2.**
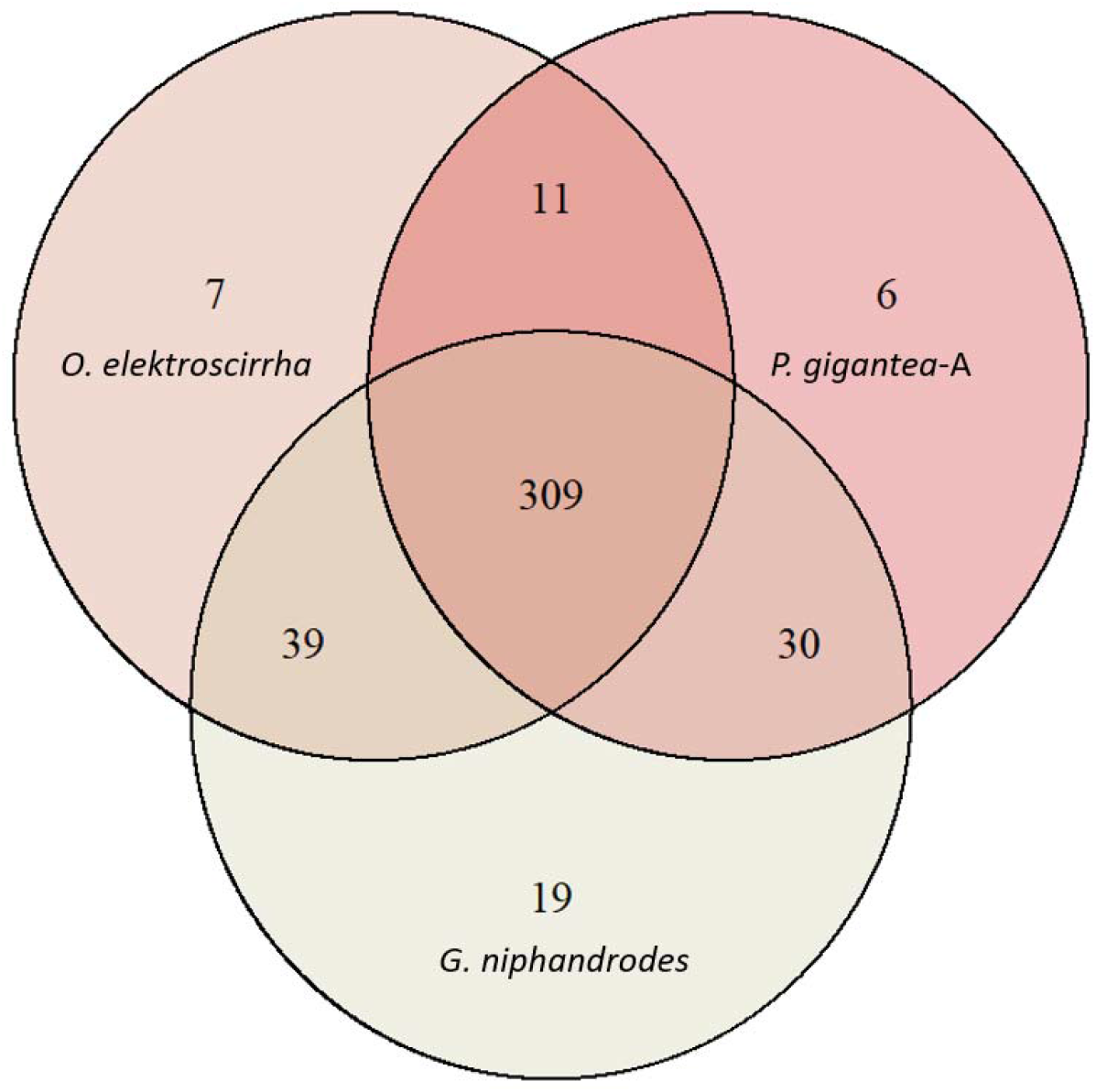
Conserved gene content overlap in sequenced gregarine Apicomplexa. Venn diagrams show the overlap in BUSCO orthologs identified from the apicomplexaodb10 dataset. Single, duplicated, and fragmented genes were all counted as present. Three distantly related gregarines all possess a large core of genes, 309 of 446 in the dataset. Moreover, *O. elektroscirrha* shares different sets of orthologs with each of the two species, suggesting independent lineage-specific gene loss across this group.

### *Genome resequencing data reveals infections in multiple* Danaus *species*

The new OE genome allows us to detect parasite DNA in short-read genomic datasets, providing a route to screen for infection from sequenced samples without needing direct access to the butterfly itself. By analysing the depth of aligned Illumina reads (with stringent filtering, see Methods) from 38 wild-collected samples representing seven *Danaus* species, we found clear evidence for infection in five out of eight *D. plexippus* from Florida and Ecuador, ten out of eighteen *D. chrysippus* from Kenya, and two out of two *D. petilia* from Australia (Figure S1, supplemental table). The remaining species were each represented by just one or two samples, so the absence of infections in these samples does not rule out that infections occur in the wild. The longest genome scaffolds provide the most robust evidence for infection. Shorter scaffolds show variable read depths in infected samples, and some have non-zero read depth in uninfected samples (Figure S1), likely indicating some shared repetitive DNA between the host and parasite genomes. Read depths tend to be low (<1X on average in most cases, compared to 10-25X coverage of host DNA), indicating that parasite DNA is far less abundant than host DNA, as expected of incidental sequencing without oocyst concentration or manual disruption.

Many of the samples considered were sequenced by other research groups, but the availability of bodies for 18 of the *D. chrysippus* samples allowed us to compare the accuracy of infection screening using genomic data versus the conventional method of microscopic detection of oocytes. This revealed nearly 100% correspondence, with nine individuals identified as infected using both methods, one with weak evidence for infection from genomic data but not from oocyst identification, and eight identified as uninfected using both methods. This implies that screening based on sequence data is at least as sensitive as the conventional approach.

### *A diverged* Ophryocystis *sequence from a related butterfly*

Using the above *O. elektroscirrha* assembly, we scanned for apicomplexan scaffolds in a previous genome assembly of a *Danaus chrysippus* sample that had tested positive for infection (sample RFK001). In total, we extracted 822 sequences, totaling 8,799,632 bases (N50 = 44,501) (Figure S2). This would account for almost an entire genome if the sequences belong to *O. elektroscirrha*, but from the start, this conspecific status was dubious. Absolute divergence between sequences was 0.05. This substantial dissimilarity between sequences motivated further analyses.

To further investigate functional divergence, we used the same GeneMark approach as above to annotate the *Ophryocystis* sequence pulled from the *D. chrysippus* assembly. An overview of this assembly is shown on the right column of Table 1. We place less emphasis on the raw gene counts and number of BUSCOs missing from this assembly, as it is more fragmented than the *O. elektroscirrha* assembly and more of it had to be filtered out for analyses, resulting in only 8.06 Mb available for gene annotation. Nonetheless, we identified 2,369 genes (2,280 protein coding) with 48.9% complete BUSCOs and 11.2% fragmented. In raw numbers, 102 orthologs were missing from this *D. chrysippus-derived* genome compared to the *D. plexippus* parasite. More intriguingly, the *D. chrysippus* parasite annotation contained 4 BUSCO orthologs that were not found in *O. elektroscirrha*. We attempted to validate these results by BLASTing the putatively missing BUSCOs in the more complete, unscaffolded assembly and found strong hits (e-value < 10^−20^) for two of the four genes. Thus, while some of these differences appear to be false negatives arising from the assembly process, there may have been independent trajectories of gene loss and retention even between these two much more closely related parasites, in addition to the broad differences between sequenced gregarines. To explore the relationship between parasite lineages from different host species, we used the *Ophryocystis*-like sequence data extracted from twelve infected butterflies to build a neighbor-joining tree. We restricted our analysis to sites at which high quality genotypes were present for at least nine of the twelve individuals. This filtered alignment included 12,976 variants across 365,363 aligned sites (4% of the genome, which is unsurprising given the low coverage and consequent high missingness). Sequences derived from *D. plexippus* butterflies (n=4) form one clade which includes the *O. elektroscirrha* reference sequence (Figure 3). Sequences from *D. chrysippus* (n=6) form a distinct clade with substantial divergence from *O. elektroscirrha* as expected based on comparison between the genome assemblies. Finally, *D. petilia* parasites (n=2) are most closely related to each other, and sister to the clade of sequences from *D. chrysippus*. This pattern of relatedness is consistent with a host specificity of parasite lineages that could lead to co-evolution and speciation.

**Figure 3.**
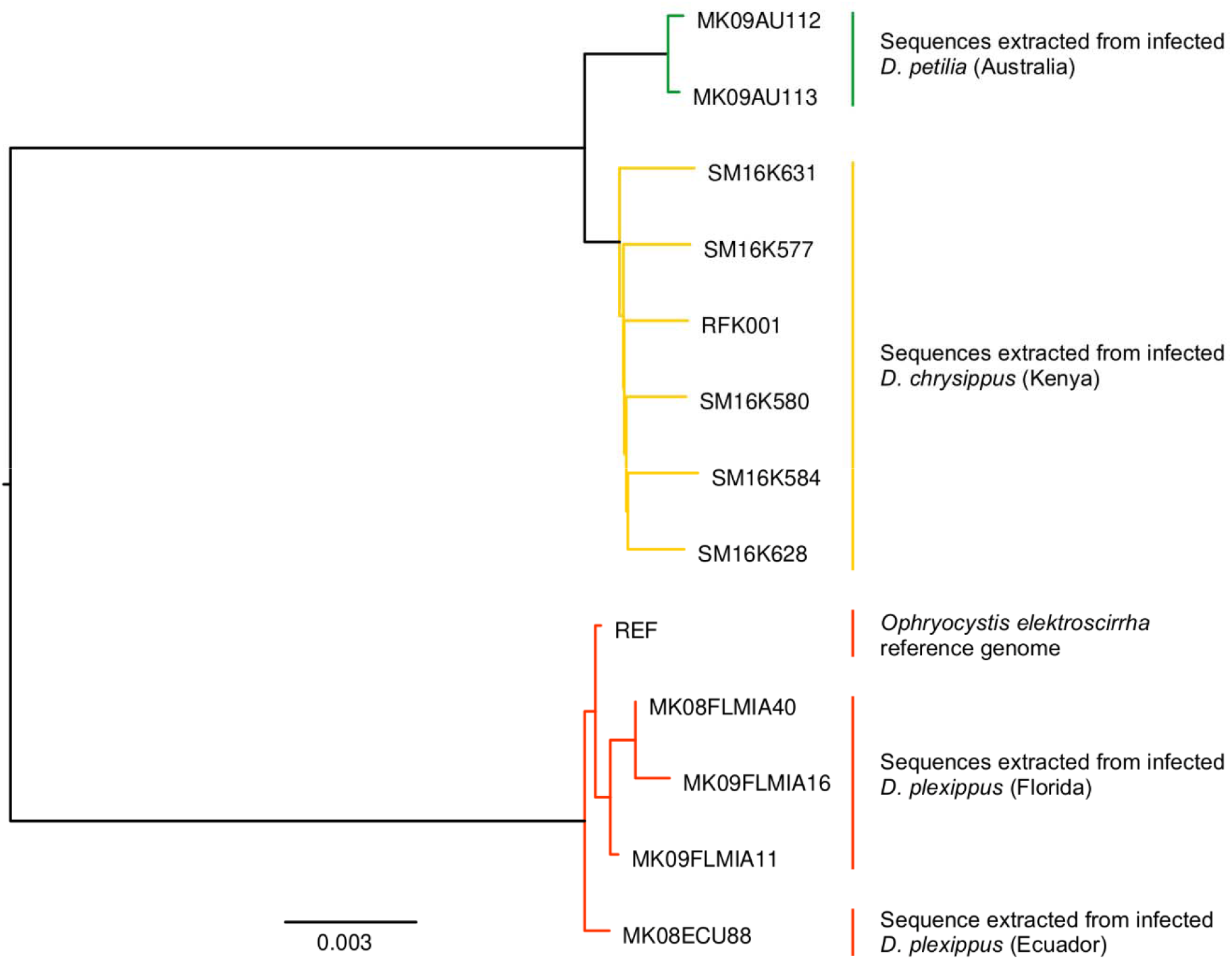
A neighbor-joining tree of *Ophryocystis* sequences pulled from the sequencing of various milkweed butterflies, including both best-studied host Danaus plexippus and other related species. Branch lengths are proportional to sequence changes. Parasite sequences cluster perfectly with host species and *Ophryocystis* samples collected from *D. plexippus* form a distinct clade from those found in other *Danaus* species.

### ATPase evolution

Using a *P. falciparum* ATPase to BLAST the annotated genes, we identified ATPases in both *Ophryocystis* assemblies as well as *P. gigantea* and *G. niphandrodes. Ophryrocystis elektroscirrha* has three ATPases identifiable in the current annotation: g016110, g003020, and g013730. Similarly, three genes are found in the *O. elektroscirrha*-like annotation: g013280, g012340, and g002420, with the latter likely fragmented as a result of the method of assembly. The other gregarines, *G. niphandrodes* and *P. gigantea* have 3 and 6 putative ATPases respectively. It is notable that the *Porospora gigantea-A* annotation shows more ATPases than the other gregarines, suggesting potential gene duplication, but given the nature of how *Porospora* was assembled, it is possible these are pooled genes from two distinct lineages (49).

We placed all of these sequences in the context of more robustly annotated ATPases using a maximum-likelihood tree of conserved amino acid subsequences of ATPases. At a coarse level, gregarines fit within established apicomplexan patterns. None of our query ATPases belong to the ENA, or Type II ATPases, which have never been reported in other Apicomplexa. Moreover, all surveyed gregarines possess SERCA and PMCA calcium ATPases and at least some species possess ATP-4 sodium ATPases (Figure 4).

**Figure 4.**
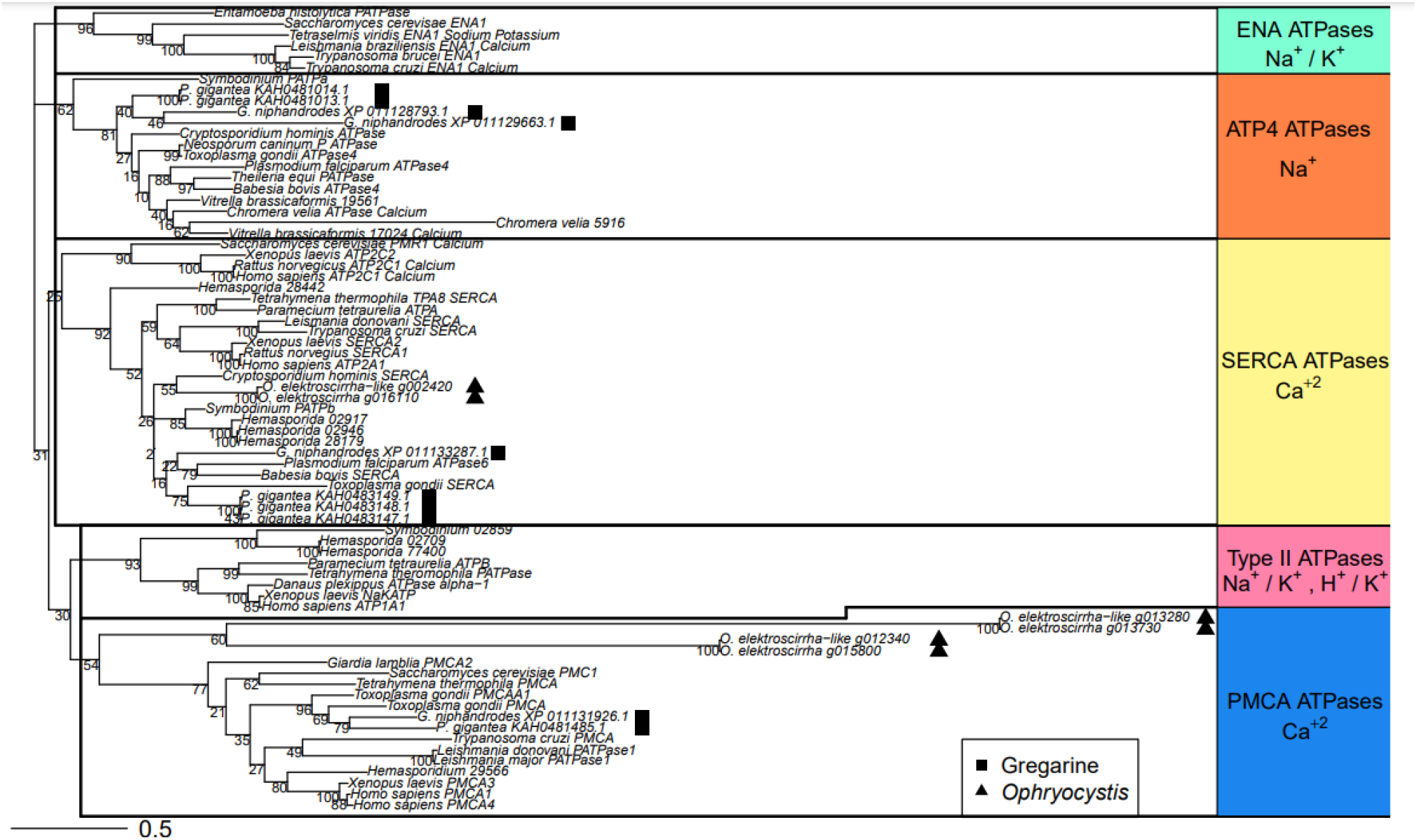
Sequence-based categorization of gregarine ATPases. We used a set of highly conserved amino acid subsequences of ATPases, adding genes from *P. gigantea, G. niphandrodes, O. elektroscirrha, O. elektroscirrha-like* to a dataset from Lehane et al. (36) and generated a maximum likelihood tree with bootstrap support for nodes. As a group, the sequenced Gregarine ATPases (squares) do not differ from other studied Apicomplexa. None of these taxa possess Type II or ENA ATPases. At least some sampled gregarines have PMCA, SERCA, and ATP-4 type ATPases (squares), but *Ophryocystis* lineages (triangles) lack this last family. Additionally, *O. elektroscirrha* and *O. elektroscirrha-like* have two PMCA-like ATPases that show substantial sequence divergence from other members of the gene family. Branch lengths are proportional to sequence changes.

The exceptions to this general pattern are more interesting, and both involve *Ophryocystis*. First, neither *O. elektroscirrha* nor *O. elektroscirrha-like* possess ATP-4 sodium ATPases. And although both lineages have a pair of ATPases that fall within the PCMA family, the branch lengths compared to the rest of the phylogeny are very long. This is not merely a matter of overall sequence divergence from other taxa, as the SERCA ATPases of *Ophryocystis* are not exceptionally different from the rest of Apicomplexa. Thus there appears to be a unique dynamic among the PMCA ATPases.

## Discussion

Here we report the assembly and gene annotation of the neogregarine parasite, *Ophryocystis elektroscirrha*. This species, along with other invertebrate pathogens, has largely been overlooked by modern genetic and genomic research. As such, the molecular characterization of this parasite has immediate value to both the broad understanding of apicomplexan biology and the specific host-parasite relationships between milkweed butterflies and *Ophryocystis*.

### Ophryocystis elektroscirrha *in relation to other Apicomplexa*

Apicomplexa are an ancient and diverse clade of eukaryotes, about which very little is known outside of human-relevant pathogens (64). As such, generating *a priori* expectations for genome size and content, as well as *a posteriori* assessment of an assembly and annotation, are difficult tasks. At the least, existing apicomplexan genomes give us some bounds for expectations. In comparison to other identified Apicomplexa, *Ophryocystis elektroscirrha* is the second smallest reported genome to-date at just under 9 megabases total, barely larger than the genomes of *Porospora* spp. (49). Similarly, it has the fewest genes, but overall gene density (genes per megabase of sequence) is in the middle of the range of studied species.

Beyond these very coarse metrics though, finding similarities in genomic organization has generally proved difficult with Apicomplexa (65), likely for both biological and methodological reasons. On the biological side, parasites often have reduced genome sizes and gene counts compared to free-living organisms, owing to both increased selection for efficiency in replication and relaxed pressures to maintain molecular mechanisms that overlap with resources found in their hosts (66,67). Indeed, such explanations have long been invoked to explain apicomplexan genome sizes and gene contents (68). Because these selective pressures are happening independently in different apicomplexan lineages, co-evolution with different hosts should result in different patterns of gene loss between parasite species. The end result is that attempting to find conserved sets of genes in the same order along chromosomes (*i*.*e*. synteny) between apicomplexan parasites has proved challenging (65).

On the methodological side, assessment of gene content is limited by available data. We recovered ∼80% of expected Apicomplexan BUSCOs (i.e., “conserved” orthologs) in our annotated gene set for *O. elektroscirrha*, a low BUSCO score by most metazoan standards. From that perspective, such a high proportion of missing orthologs could be indicative of an incomplete or incompletely annotated assembly; however, the set of genes considered as “universal single copy” for a clade of organisms is defined only by existing genetic data (69). Thus, genes that appear universally conserved within the small handful of well-characterized Apicomplexa may be truly lost in the *Ophryocystis* lineage. Indeed, a recent study of another gregarine parasite genus, *Porospora*, generated assemblies for two species, both of which with roughly ∼60% of expected apicomplexan BUSCOs; a broader comparison in that same study identified 83% of the expected BUSCOs in *Gregarina niphandrodes* (49). In that context, the conserved gene content of *O. elektroscirrha* fits well within gregarine expectations.

When we directly compared the overlap of BUSCOs identified in *Porospora* and *G. niphandrodes* to *O. elektroscirrha*, found that only a small fraction (∼6%) were truly absent from all three. More commonly, genes were conserved in only one or two species (26%). *Ophryocystis elektroscirrha*, despite having roughly half the annotated genes as the other two genera, was not merely a subset of either species in gene content. Instead, it displayed independent overlaps with the other two, as expected if co-evolution with different hosts and environments has driven unique patterns of gene loss in each lineage. More generally, these results suggest that the gregarine apicomplexans have a smaller set of conserved genes than currently recognized for the better-studied clades of Apicomplexa.

### Cryptic diversity of milkweed butterfly parasites

Using the *O. elektroscirrha* genome, we were able to recover an apicomplexan genome from another milkweed butterfly, *Danaus chrysippus*. This sequence was highly diverged from the *O. elektroscirrha* genome, with absolute sequence divergence at roughly 5%. This number, which is comparable to the level of divergence between the two host butterfly species, may even be an underestimate, owing to our method of sequence discovery. Because we used similarity to the *O. elektroscirrha* genome to find new sequences, any regions that are exceptionally quickly evolving would not be detected via this method. Nevertheless, the comparable genome size suggests we are capturing the vast majority if not the entire parasite sequence.

We took these related *Ophryocystis* sequences through the same gene annotation pipeline we used for *O. elektroscirrha*. These new sequences were more fragmented, and thus unsurprisingly, we annotated fewer genes from the *D. chrysippus* derived parasite than from the *O. elektroscirrha* assembly. As such, apparent absences from the *chrysippus-*derived genome may be false negatives. In the other direction however, two apicomplexan BUSCOs were identified in the *chrysippus* parasite sequences that were absent from the *O. elektroscirrha* annotation. It is possible that these also represent false negatives, since both genomes were trimmed of short sequences prior to annotation; however, it is not outside the realm of possibility that these lineages may have different constraints on gene content if they consistently associate with different host species.

To explore the host and parasite relationship, we expanded sampling to sequenced reads from multiple infected *D. plexippus* and *D. chrysippus* and created a phylogenetic tree to examine their relatedness. We recovered a pattern of reciprocal monophyly for parasites based on host species. This pattern, along with the sequence and potential gene content divergence all suggest that different species of host may harbor distinct species or at least substantially differentiated lineages of *Ophryocystis*.

Work using only the 18S ribosomal RNA sequence found that parasites isolated from *Danaus plexippus* and the distantly-related moth *Helicoverpa amigera* clustered in a similar pattern (31), but could not explore this pattern at larger scale without genomic data. More to the point, it is less surprising to see differentiation between parasites of hosts separated by ∼110 million years of evolution (70) than to see such a pattern within a single host genus. It raises the possibility that other species of *Danaus* with a reported apicomplexan parasite are infected by distinct species of *Ophryocystis*.

Indeed, earlier experimental evidence has hinted at such a possibility. In cross-infection experiments exposing *D. plexippus* and *D. gilippus* hosts to *Ophryocystis* collected from either an intra- or interspecific source, the parasites were most successful infecting the same species of host from which they were first collected (29). In other words, *Ophryocystis* exhibits significant host-specificity. What remains to be seen is how *Ophryocystis* lineages have evolved with *Danaus*. It may be that host and parasite share very similar speciation histories, as seen in other systems (e.g. birds and lice: Hughes *et al*. 2007). Alternatively, given that many species of milkweed butterfly share the same host plants in sympatric ranges, host switching may be driven more by an ecology of opportunity than phylogenetic history.

### Milkweed, butterflies, parasites, and ATPases

The interactions between milkweed-feeding insects and their food source chemistry are well-studied. Milkweed toxicity derives in large part from a class of cardiac glycoside compounds that bind to and inhibit sodium potassium pump (Na ^+^ /K ^+^ ATPase) proteins of the animals that ingest them (72). For multicellular animals, sodium potassium pumps are key to the process of establishing an ion gradient across cell membranes and in particular allow for proper transmission of electrical signaling at the intercellular level. To maintain this crucial function, many milkweed-feeding insects have evolved a small set of amino acid substitutions in their ATPase sequence that confer resistance to cardiac glycoside binding. Unrelated herbivores such as butterflies and beetles show two convergent changes, a valine and histidine substitution in the α subunit of their Type II ATPase (26,32). Even more distantly related taxa, including a wasp parasitoid, a nematode parasite, and a bird predator of monarchs, all have similar substitutions in their ATPases (34). Together these results suggest that a consistent selective pressure has driven a convergent molecular solution in independent lineages. What remains to be seen is if similar dynamics have occurred in non-metazoan members of the milkweed community. *Ophryocystis* parasites of milkweed butterflies spend much of their life cycle in the larval gut or other host tissues (8) and are routinely exposed to cardiac glycosides. Experimental evidence shows that *O. elektroscirrha* growth is negatively impacted by the presence and concentration of cardiac glycosides (27) and that infected female *D. plexippus* (which would likely transmit *O. elektroscirrha* to their offspring) preferentially choose to lay eggs on milkweed with more cardiac glycosides when given a choice (73). Thus, the hosts and their ingested phytochemicals appear to exert a selective pressure on the parasite, but what their molecular targets are had yet to be explored.

Aside from a lack of sequence data, the main challenge to this line of research is that between Metazoa and Apicomplexa there is a lack of homology, in both protein sequence and function. First, although ATPases are an evolutionarily ancient class of proteins found in protists as well, they obviously cannot play roles in intercellular electrical signaling of a single-celled organism, so the mechanism of toxicity cannot be the same. But ATPases are still important regulators of cellular homeostasis with respect to salt and pH balance and have been suggested as targets for drug development (37,74). Even if their functions are different, inhibition of apicomplexan ATPases would still be detrimental to the organism.

In the above descriptions of ATPase – cardiac glycoside interactions in Metazoa, the target ATPase is always the Type II Na^+^/K^+^ ATPase. No Apicomplexa are known to possess this specific family of ATPase and, until recently, they were thought to lack any sort of Na^+^ ATPase. More recently however, it has been shown in *Toxoplasma gondii* that the ATP-4 ATPase, previously thought to employ calcium, is in fact a sodium ATPase (36). This family could be a candidate for the cardiac glycosides’ target, given the conserved cation use. However, none of the ATPases had been characterized in gregarines.

We used conserved sequence domains across ATPases to characterize the specific families of ATPases present in *P. gigantea* and *G. niphandrodes, O. elektroscirrha*, and *O. elektroscirrha-like*. As a whole, the gregarines fit with better-studied Apicomplexa. None possess Type II or ENA ATPases, and all possess PMCA and SERCA calcium ATPases. Intriguingly though, both *Ophryocystis* lineages apparently lack ATP- 4 sodium ATPases that are found in *Porospora* and *Gregarina*. As the only putative sodium ATPase in Apicomplexa, is tempting to speculate that loss of this ATPase may have been related to cardiac glycoside presence in the host. A parasite ATPase that is routinely inhibited by the host’s chemical environment is essentially non-functional and may be lost under mutation accumulation. Of course, this line of reasoning relies on only a small set of observations; it would be bolstered by study of closer relatives to *Ophryocystis* that parasitize non-milkweed-feeding insects.

Considering the ATPases still present in *Ophryocystis*, we recovered three putative calcium pumps, a SERCA (localized to the endoplasmic reticulum within the cell) and two PMCA ATPases which are located on the plasma membrane of the cell. These should be considered as potential targets of cardiac glycosides, as some sodium pump blockers may have non-specific inhibitory action against calcium pump ATPases as well (75). The PMCAs would be most likely to interact with cardiac glycosides, which canonically affect plasma membrane proteins (76). In *Ophryocystis*, these proteins’ sequences are very different from other sequenced Apicomplexa; in contrast, the SERCA ATPases of *Ophryocystis* do not show exceptional sequence divergence from orthologous genes. Thus, only the plasma membrane associated ATPases of *Ophryocystis* appear highly diverged. It will require more gregarine sequences for comparison, but these proteins are intriguing candidates for the target of cardiac glycoside toxicity in milkweed butterfly parasites.

## Conclusions

Sequencing the genome of *Ophryocystis elektroscirrha* has yielded immediate insights into potential biochemical evolution driven by milkweed chemistry, cryptic variation in milkweed butterfly parasites, and the true extent of gene turnover across Apicomplexa. We present these results to facilitate further exploration of these parasites to give context to the large body of disease ecology research on this clade. We hope that our novel data collection methods, both in DNA extraction and parasite sequence screening, will aid in future work on poorly understood Apicomplexa.

## Data Availability

The genome assembly for *Ophryocystis elektroscirrha* can be found with the accession JAQIFP000000000. Raw DNA reads used to assemble the genome and RNAseq used to scaffold can be found with PRJNA906508. The annotation is included in the Supporting Materials. The *O. elektroscirrha-like* assembly and annotation can also be found as supplementary material; the assembly in particular we chose not to formally archived due to its potentially incomplete nature an uncertain taxonomic status. Accessions for the butterfly sequences used to screen for *Ophryocystis* reads can be found in the Supplement as well.

## Supporting information

Supplemental table of accessions and infection screening

## Acknowledgements

The authors wish to thank Adele Lehane, Adelaide Dennis, and David Heckel for insights into ATPase evolution; Sonia Altizer and Maria Luisa Muller Theissen for discussions about OE – butterfly interactions; and Jasmin Albert, Thomas Johnson, and Megan Hansen for additional help in protocol optimization of DNA extraction. James Walters was supported by the US National Science Foundation grant NSF-ABI 1661454. Rachel Manweiler and Jordyn Koehn were supported by the Gould Summer Entomology Fellowship from the University of Kansas

