## Supplementary Figures for "Genome sequence of *Ophryocystis elektroscirrha*, an apicomplexan parasite of monarch butterflies: cryptic diversity and response to host-sequestered plant chemicals"

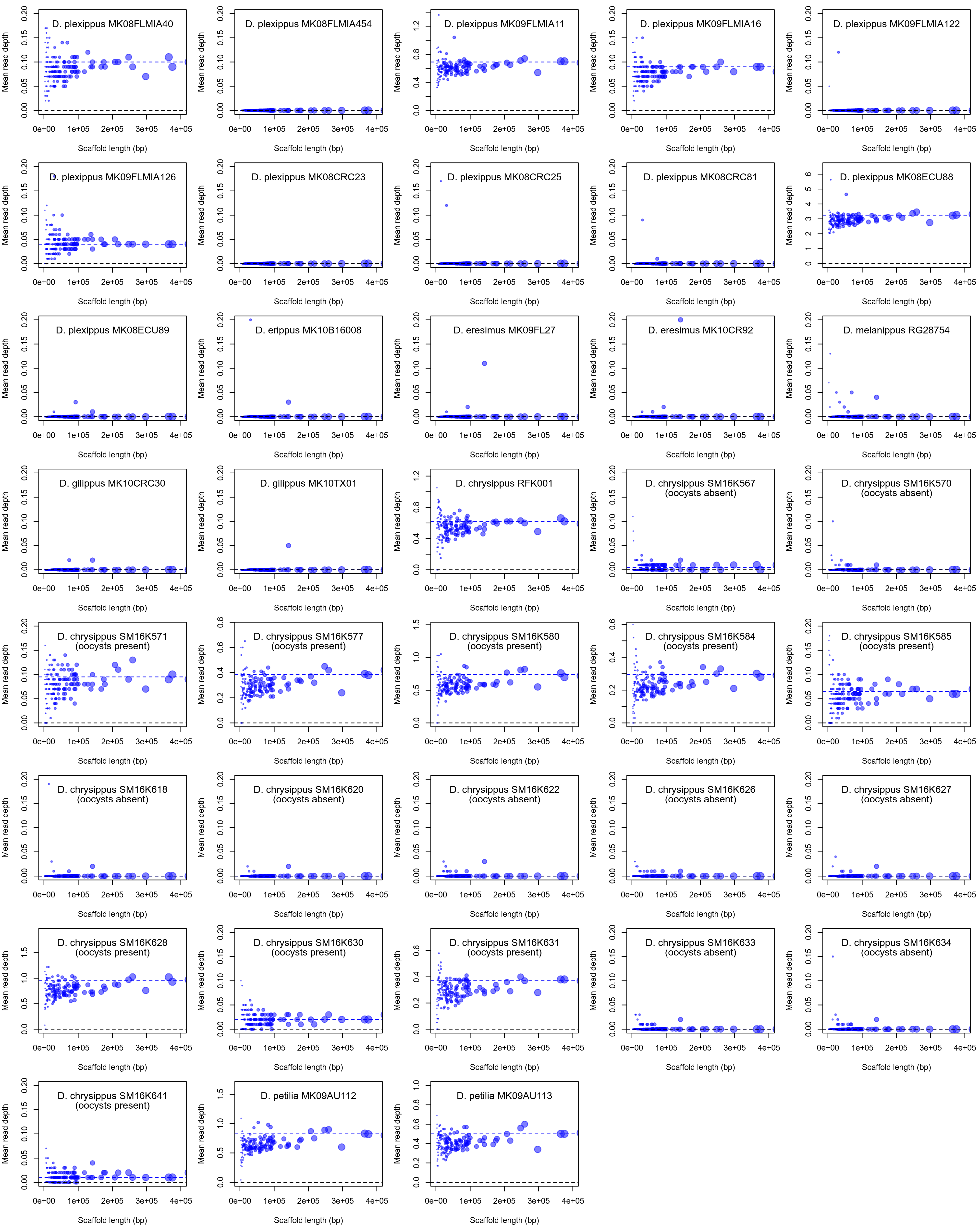

**Figure S1. Using genomic data from butterflies to diagnose *Ophryocystis* infection**

Mean read depth per scaffold in the *O. elektroscirrha* genome for Illumina sequencing reads generated from butterfly tissue is plotted against scaffold length. Each plot represents a separate butterfly sample, with species and ID indicated. Although short scaffolds show variable read depth, longer scaffolds can reliably detect infection in the form on non-zero average read length. Dashed horizontal lines indicate the median read depth for scaffolds >200kb in length. For 18 *D. chrysippus* samples, it is also indicated whether oocytes were detected on the sample. All nine positive cases correspond to cases where read depth is also non zero.

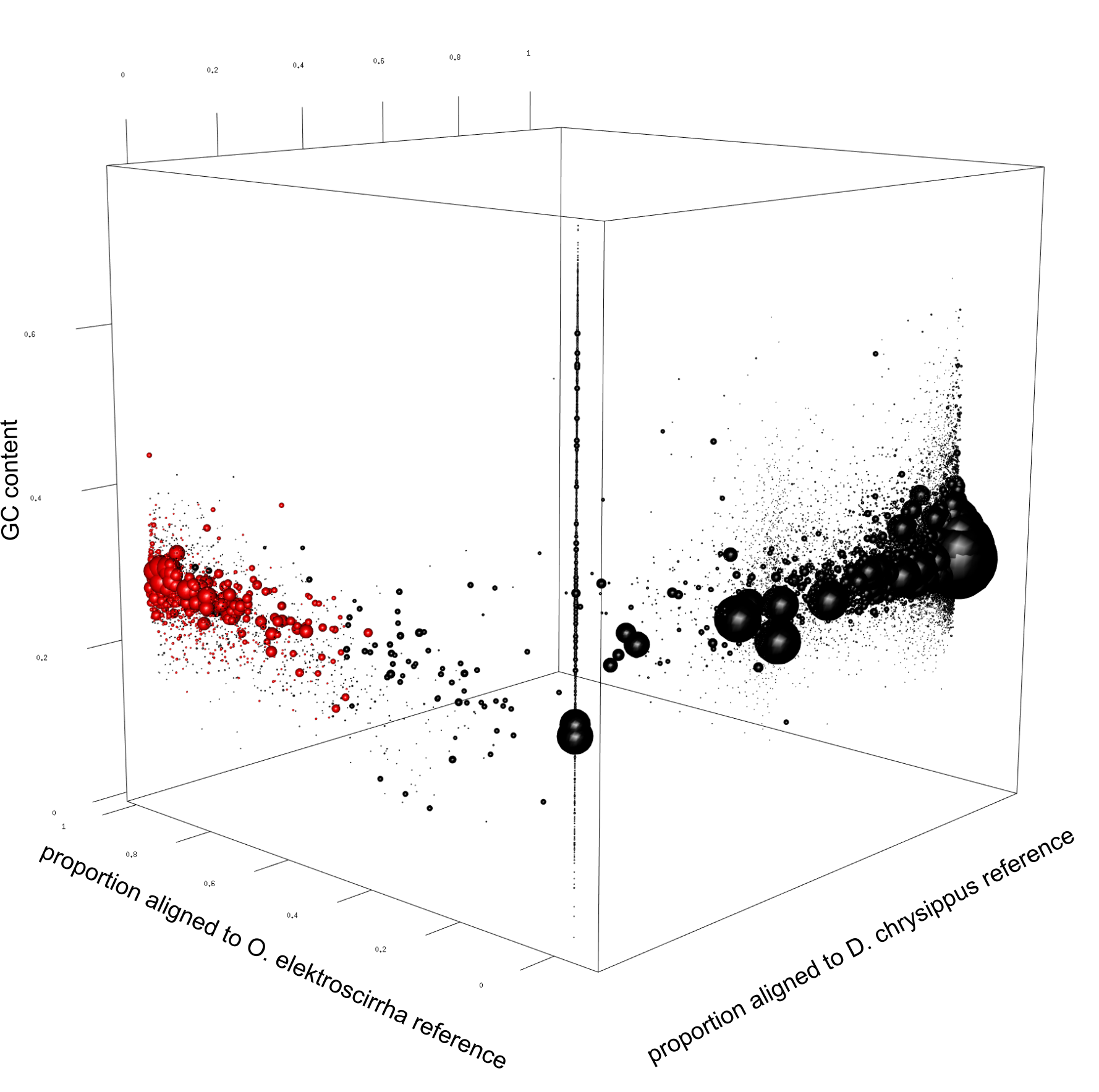

**Figure S2. Identification of *Ophryocystis*-like scaffolds in a *D. chrysippus* genome assembly**

Scaffolds identified as sufficiently *Ophryocystis*-like to represent a putative genome for an *Ophryocystis*-like parasite of *D. chrysippus* are indicated in red. GC content (vertical axis) is only slightly different on average between the host and parasite genomes, so this was not considered for identification of parasite scaffolds.

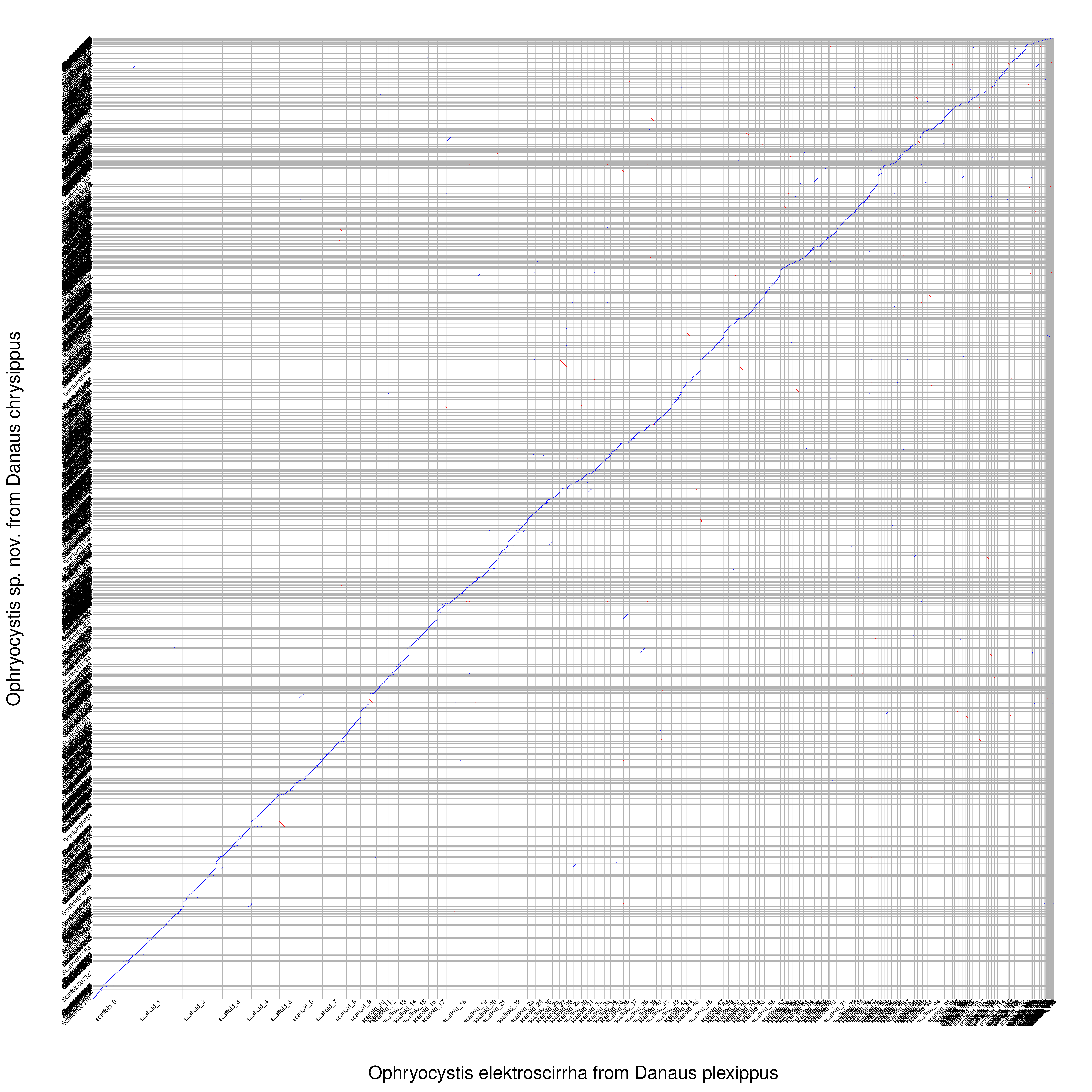

**Figure S3. Alignment between the genomes of O. *elektroscirrha* and the *Ophryocystis*-like parasite of *D. chrysippus***

Blue diagonal lines indicate tracts of aligned sequence between the two genomes. Scaffolds are separated by gray lines. The fact that most scaffolds have a complete and 1:1 alignment indicates that the genomes are likely both close to complete.
